# Label-free Cell Tracking Enables Collective Motion Phenotyping in Epithelial Monolayers

**DOI:** 10.1101/2021.12.14.472148

**Authors:** Shuyao Gu, Rachel M. Lee, Zackery Benson, Chenyi Ling, Michele I. Vitolo, Stuart S. Martin, Joe Chalfoun, Wolfgang Losert

## Abstract

Collective cell migration is an umbrella term for a rich variety of cell behaviors, whose distinct character is essential for biological function, notably for cancer metastasis. One essential feature of collective behavior is the motion of cells relative to their immediate neighbors. We introduce an AI-based pipeline to segment and track cell nuclei from phase contrast images. Nuclei segmentation is based on a U-Net convolutional neural network trained on images with nucleus staining. Tracking, based on the Crocker-Grier algorithm, quantifies nuclei movement and allows for robust downstream analysis of collective motion. Since the AI algorithm required no new training data, our approach promises to be applicable to and yield new insights for vast libraries of existing collective motion images. In a systematic analysis of a cell line panel with oncogenic mutations, we find that the collective rearrangement metric, D^2^_min,_ which reflects non-affine motion, shows promise as an indicator of metastatic potential.

**Graphical Abstract:** 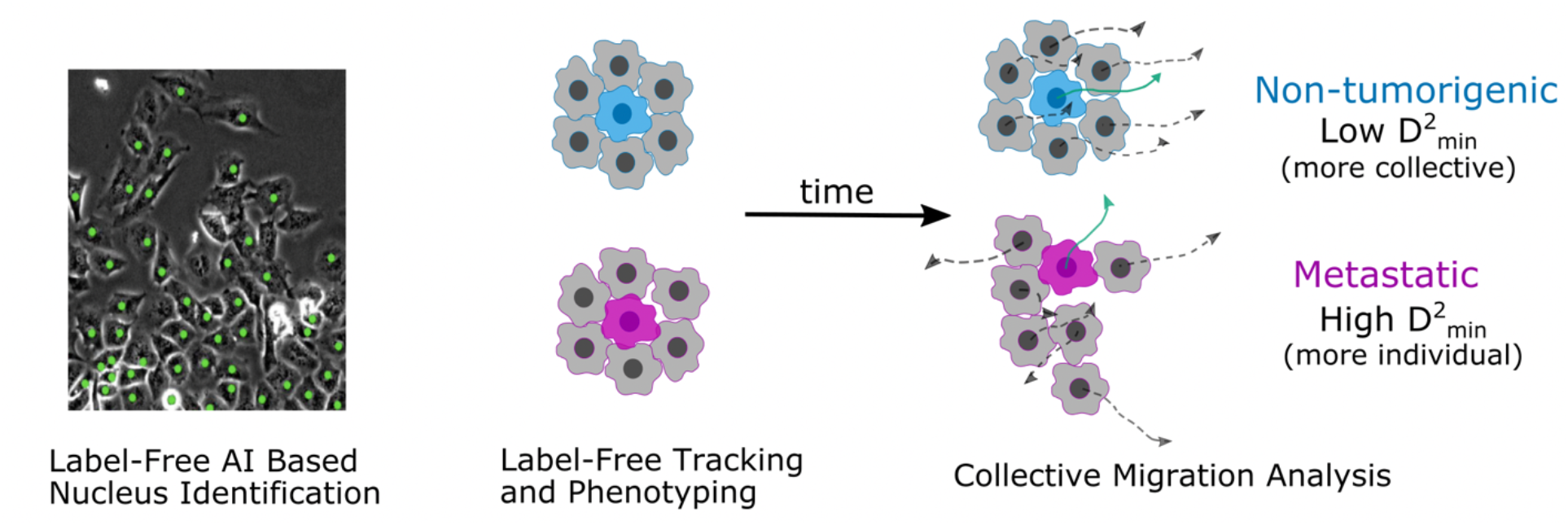

**Highlights:**

- Versatile AI-based algorithm can robustly identify individual cells and track their motion from phase contrast images.
- Analysis of motion of cells relative to nearby neighbors distinguishes weakly tumorigenic (KRas) and metastatic (KRas/PTEN^*-/-*^) cell lines.

## Introduction

Breast cancer is one of the most common cancers in women. Many breast-cancer-related deaths result from metastasis, a complex, multistep process in which cancer cells migrate away from the primary tumor and eventually grow in other organs [1-3]. During the metastatic process, clusters of primary tumor cells can leave the original site, circulate in the bloodstream, and seed secondary tumor growth at distant sites. There is growing evidence that tumor cells that collectively migrate and enter the bloodstream as clusters (defined as any group larger than one cell) are 50-100 times more likely than individual cells to lead to metastasis [4-6]. Using phase contrast movies of migrating cells, we introduce a new Artificial Intelligence (AI) based approach to characterize a shifting balance between individual motion and collective motility of mammary epithelial cells as they become more transformed, tumorigenic, and metastatic.

We use a commonly studied breast epithelial cell line (MCF10A), a highly metastatic cell line (MDA-MB-231), and a genetically defined model system for cancer and metastasis. The genetically defined model system is based on the MCF10A cell line and introduces phosphatase and tensin homolog deletion (PTEN^-/-^) and/or overexpression of activated KRas(G12V), two common oncogenic mutations that have downstream impacts on the cytoskeleton. PTEN loss is found in 24% of breast cancers [7] and has been associated with poorer prognoses in many solid tumors [8]; KRas is a frequent driver of tumors across different types of human cancers [9, 10]. This model cell system relates our results to *in vivo* outcomes, since the cell lines have been characterized in terms of tumorgenicity [11] and metastatic behavior [12].

In a previous study, we imaged the motion of groups of cells from these cell lines in a well-defined in-vitro assay [13] and characterized the motion using particle image velocimetry (PIV). We found that deletion of PTEN caused cells to move more collectively, while KRas activation had the opposing effect [14]. However, when both mutations were present, the phenotype induced by KRas dominated and the KRas and KRas/PTEN^-/-^ cell lines displayed similar dynamics [12, 14], even though they have significantly different in vivo outcomes. One limitation of PIV is that it characterizes motion based on the movement of all features in an image. Thus, PIV captures overall movement visible in an image, including the motion of the many organelles visible in a typical phase contrast image, which may obscure the full motion of each cell’s nucleus. However, the cell nucleus serves as the main reference point of a cell, steering cell motion [15] and defining orientation when compared to the microtubule organizing center [16].

In this study, we introduce an AI-enabled (U-Net) cell nucleus segmentation tool for epithelial cells [17] that allows us to track cell nuclei and study their dynamics. The AI model was trained on stem cell images with Lamin B1 stain for automated nucleus detection [17] and did not require further training to perform nucleus segmentation on the studied epithelial cell sheets. Additionally, our AI-enabled single-cell tracking allows us to characterize motion of cells relative to their neighbors. In particular, we calculate D^2^_min_, which describes the extent of similarity of motion between the cell of interest and its closest neighbors [18, 19]. This ability to assess how closely a cell follows its neighbors yields a phenotype of cell rearrangements during collective migration capable of differentiating the metastatic KRas/PTEN^-/-^ from the non-metastatic cell lines. AI-enabled tracking and quantification of collective rearrangements thus provide new tools for quantifying metastatic potential and could be used to improve patient diagnosis or to develop novel therapeutic targets.

## Results

### AI-based analysis workflow enables label-free single cell tracking

Our analysis focuses on the collective dynamics of a migrating monolayer of epithelial cells (**Figure 1a)** imaged with phase contrast microscopy. Nuclear segmentation and tracking are a powerful approach to studying cell migration dynamics. Here we show that deep learning (U-Net) based inferences of nuclear positions can provide a substitute for fluorescently labeled nuclei. We use a successful image analysis deep learning model, U-Net, that was originally trained on stem cell (a human iPSC clonal line) phase contrast images and ground truth nucleus data (**Figure 1b**) [17]. We find that the model can identify the locations of cell nuclei from only phase contrast images of other mammary epithelial cell lines, such as the MCF10A epithelial cell line (**Figure 1c**), without requiring any training on these epithelial cell lines. Once segmented, the images are fed into a tracking algorithm [20], which identifies a center for each segmented nucleus and maps these positions from frame to frame into a cellular track, allowing us to follow individual cell nuclei over hundreds of frames (**Figure 1d**).

**Figure 1:**
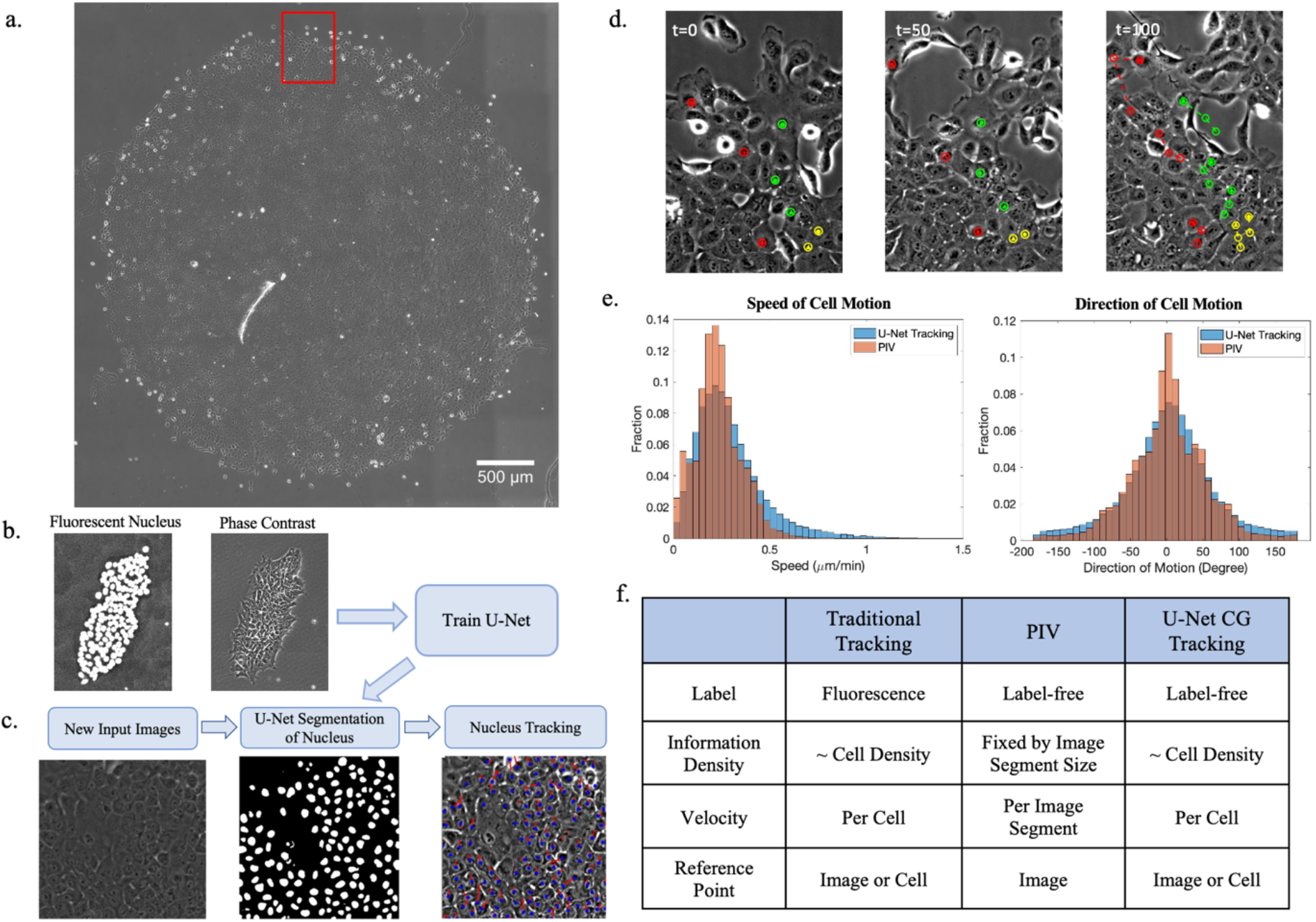
AI-based segmentation and tracking workflow captures collective migration behavior. U-Net enabled tracking of cells (a) A tiled image of a migrating cell sheet. The red box indicates the size of the field of view captured over time. (b) An example of human iPSC clonal line images used as the training set for U-Net. (c) Computational workflow: an input image is segmented using U-Net and then the centroids of the segmented cell nuclei are tracked using the Crocker-Grier algorithm. (d) Tracks of 8 cells from t = 0 to frame 100 (3 minutes per frame, frames t = 0, 50, and 100 shown) (e) Comparisons of tracking results with PIV results. (f) Table comparing benefits of traditional tracking, PIV, and U-Net Crocker-Grier tracking.

The trajectories of cell nuclei using this approach are validated by a comparison to the dynamics obtained in prior studies on the same data using particle image velocimetry (PIV) analysis [14] as shown in **Figure 1e**. The two methods produce similar speed and direction of motion distributions, with our U-Net-based tracking exhibiting longer tails at higher velocities and larger angles. Bhattacharyya coefficients quantify the overlap between distributions and range between 0 (no overlap) and 1 (complete overlap). The distributions shown in **Figure 1e** have Bhattacharyya coefficients of 0.971 (speed) and 0.991 (direction), which indicates agreement between the PIV and U-Net results [21]. We note that the angles are computed relative to the direction of monolayer expansion. The small differences observed in the tails of the distributions agrees with a previous comparison of PIV and tracking [13], which found that the motion captured by PIV is smoother than the motion of nuclei.

Our U-Net based cell tracking approach from phase contrast images complements both PIV and traditional fluorescent nucleus tracking as illustrated in a comparison table (**Figure 1f**). PIV is widely used because it can be applied to label-free images, such as the phase-contrast images studied in this current work. Fluorescent labeling of cell nuclei can provide additional information about individual cell behavior but often suffers from issues of phototoxicity [22]. Our label-free U-Net cell tracking approach yields the best of both worlds, because it allows for nuclear tracking without interfering with long-term behavior or decreasing the survival rate of cells.

### A quantitative metric for cell rearrangements distinguishes tumorigenic and metastatic cell lines

With the ability to follow individual cells in a sheet we can compute how cells move relative to their neighbors. One measure of collective rearrangements, D^2^_min_ (**Eq 1**), has been used in models of elastoplastic deformation of amorphous solids or granular materials [23] to describe the motion of particles relative to their neighbors. Analyzing a cluster of N particles, D^2^_min_ characterizes departures from the local affine deformation by fitting overall motion of the cluster over a time Δ*t* to an affine deformation and reporting the residue.

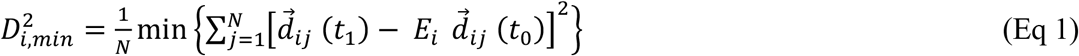

In **Eq 1**, 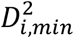 is the D^2^_min_ value for cell *i* at time *t*_1_ = *t*_0_ + Δ*t* ; N is the number of neighboring cells for cell 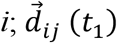 and 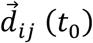 are the relative spacings between cell *i* and neighboring cell j at times *t*_1_ and *t*_0_ respectively; and *E*_*i*_ is the strain tensor for the neighbor cell group around cell *i*. We chose N= 6 under the assumption that epithelial cells have on average six neighbors in a confluent sheet. D^2^_min_ [18, 19] (**Eq 1**) is a least-squared calculation in which the final relative spacing between cells, *d*_*ij*_(*t*_1_), is predicted via a linear transformation, *E*_*i*_, of the initial relative spacing *d*_*ij*_(*t*_0_), (**Figure 2a**). The resulting difference is called non-affine (non-linear) motion, whereas the *E*_*i*_*d*_*ij*_(*t*_0_) is the affine motion. *E*_*i*_ is fit to minimize the difference between the affine motion and the actual motion. The affine component, which reflects collective motion, is essentially how well the displacements of the neighboring cells can predict the motion of the central cell. A lower D^2^_min_ value indicates the cell motion is more affine, meaning the cell more closely follows its neighbors’ motion and is moving more collectively. A higher value of D^2^_min_ thus implies a more individualistic or non-affine motion.

**Figure 2:**
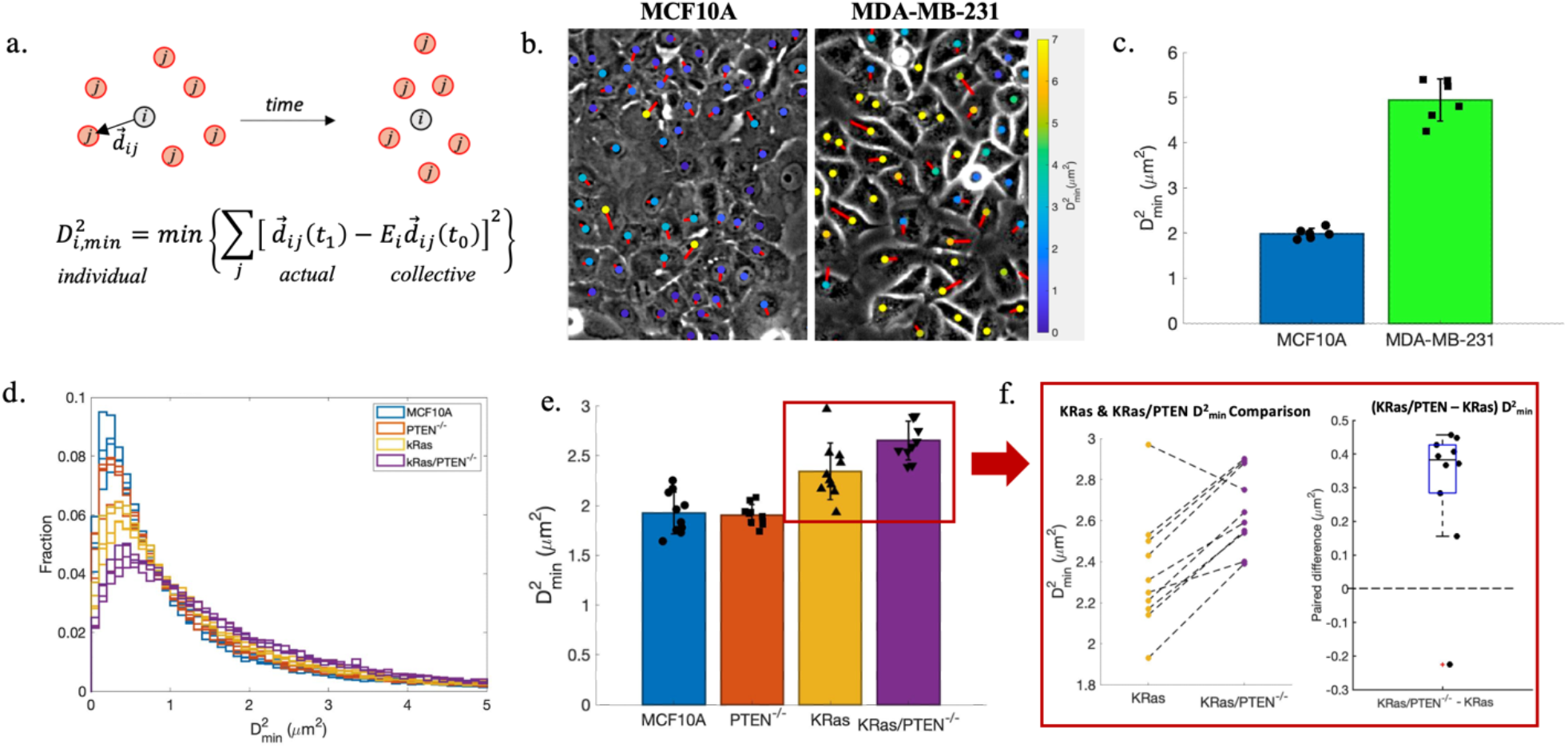
Cell rearrangements distinguish tumorigenic and metastatic cell lines. (a) A schematic of how D^2^_min_ is calculated. (b) Sheets of migrating cells labeled by their motion and rearrangements. Colorful dots represent D^2^_min_ values for individual cells, with their values indicated by the color bar. Red arrows show the direction of cell motion. (c) Comparison of average D^2^_min_ values for MCF10A and MDA-MB-231 cells over 6 independent experiments. (d) Comparison of D^2^_min_ distribution of MCF10A mutants in one set of experiments (4 technical replicates of each mutant). (e) Average D^2^_min_ for each mutant over 10 independent experiments. (f) Pair-wise comparison of KRas and KRas/PTEN^-/-^ D^2^_min._ (left) Dashed lines connect experiments performed on the same 12 well plate. (right) The central line indicates the median value of the paired difference, while the top and bottom edges of the box indicate the 75th and 25th percentiles. Error bars in c, e represent the standard deviation across independent replicates.

Using our proposed U-Net based segmentation and tracking workflow, we calculate D^2^_min_ for both MCF10A (non-tumorigenic) and MDA-MB-231 (metastatic) breast cells. This reveals strong differences in their rearrangement behavior. **Figure 2b** shows single frames from a time-lapse series in which cell nuclei are marked using colorful dots according to their respective D^2^_min_ value. The non-tumorigenic MCF10A cells behave more collectively (lower D^2^_min_ values) compared to the metastatic MDA-MB-231 cells (**Figure 2c**).

We also applied our U-Net cell tracking workflow, without additional training of the AI algorithm, on a genetically defined cancer progression model system based on the MCF10A cells [11] (**Supplementary Video S1**). In this model system, cells with PTEN deleted (PTEN^-/-^) have previously been found to remain dormant *in vivo*. KRas activated cells also usually stay dormant *in vivo*, however, sometimes the KRas cells can form primary tumors (poorly tumorigenic) [11, 14]. The double mutant (KRas/PTEN^-/-^) is aggressively tumorigenic [11] and metastatic [12]. **Figure 2d** showcases the D^2^_min_ distribution from four technical replicates for each cell line from one experiment. The D^2^_min_ distribution for each cell line is consistent across technical replicates and the combined experiment shows the overall trend that the original MCF10A and PTEN^-/-^ mutants have smaller D^2^_min_ values, which means that their motion is more collective, and cells tend to follow their neighbors’ motion. The overall results from 10 independent experiments (**Figure 2e**) confirm the poorly tumorigenic KRas and metastatic KRas/PTEN^-/-^ mutants have higher D^2^_min_ than the non-tumorigenic MCF10A and dormant PTEN^-/-^ mutants. To further differentiate the KRas and KRas/PTEN^-/-^ mutants, a pair-wise comparison was performed (**Figure 2f)**. A blocked experimental design allows us to pairwise compare the D^2^_min_ value within each experiment, depicted by dashed line connecting the raw values (**Figure 2f**, left), and by directly subtracting the KRas D^2^_min_ value from the KRas/PTEN^-/-^ D^2^_min_ value (**Figure 2f**, right). The results show that most experiments yield the metastatic KRas/PTEN^-/-^ having higher D^2^_min_ than the poorly tumorigenic KRas mutant. Therefore, the KRas/PTEN^-/-^ mutant motion is more individual (non-affine) than the KRas mutant, and D^2^_min_ can distinguish the poorly tumorigenic KRas and metastatic KRas/PTEN^-/-^ cell lines.

### Affine motion quantifies changes in collective behavior

To further study the collective component of cell motion, we extract affine dilation and deformation components from the minimized linear transformation matrix *E*_*i*_ (**Eq 1**) [18]. We decompose the 2D matrix *E*_*i*_ into a product of two matrices: a pure rotation matrix *R(θ)* and a symmetric strain tensor *F*. The average of the eigenvalues of the strain tensor F correspond to affine dilation, while the difference is the affine deformation. Physically, the two eigenvalues describe the contraction or expansion in the direction of their respective eigenvectors. Affine dilation represents the collective contraction or expansion of the group of 6 neighbors. Affine deformation describes whether the cells’ neighbors are compressed or stretched. Schematics shown in **Figure 3a** illustrate examples of affine dilation and deformation in a 6-neighbor system.

**Figure 3:**
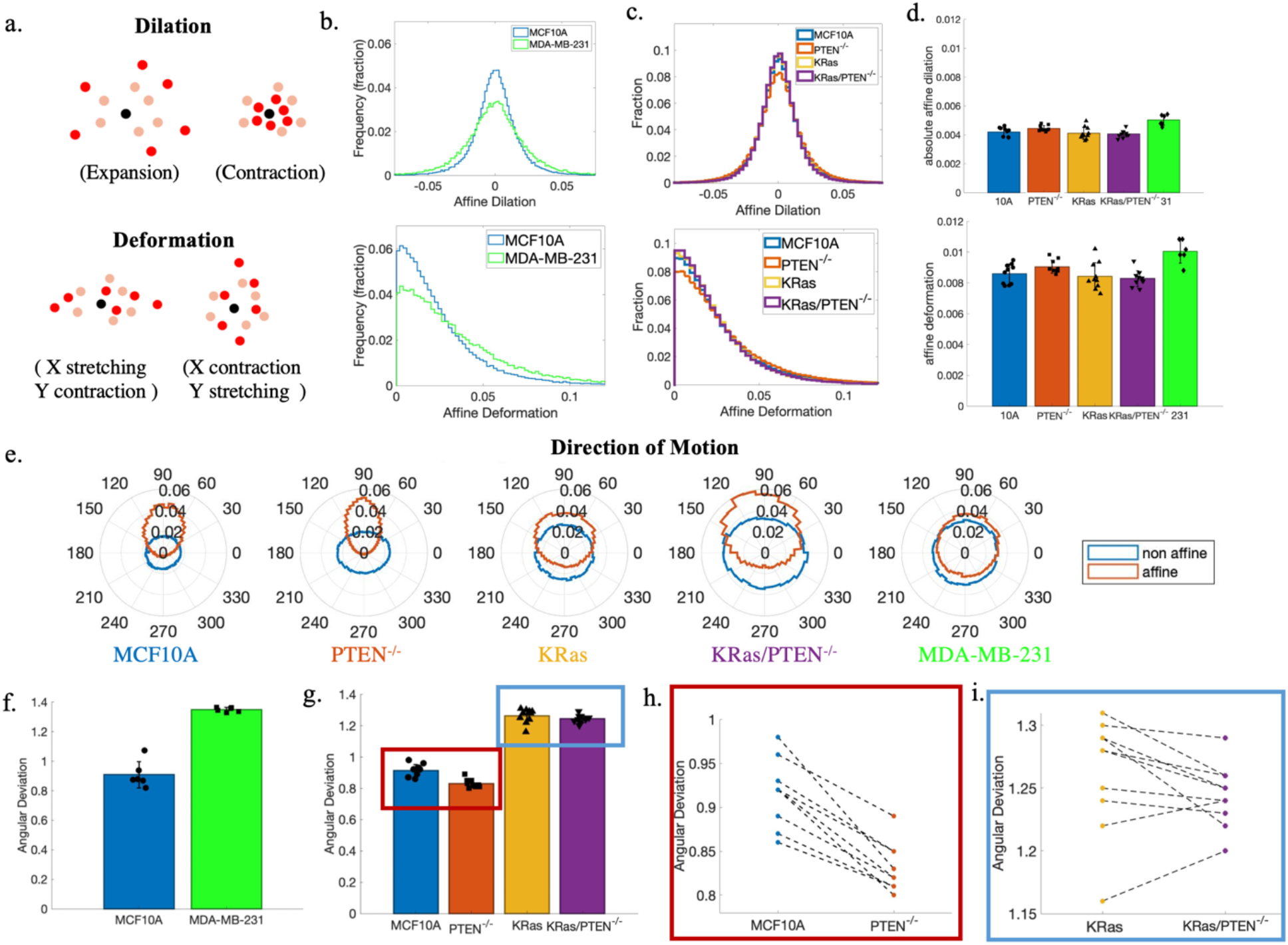
The affine component of motion describes collective behavior. (a) Schematics of affine dilation (top) and affine deformation (bottom). (b) Representative distributions of affine deformation and dilation comparing the MCF10A and MDA-MB-231 cells. (c) Representative distributions of affine deformation and dilation comparison for the MCF10A mutants. (d) The absolute value of affine dilation and average affine deformation for MCF10A mutants (10 independent experiments) and 231 cells (6 independent experiments). (e) The direction of affine and non-affine motion for representative experiments for the MCF10A mutants and MDA-MB-231. (f) MCF10A and MDA-MB-231 affine angular deviation. (g) MCF10A mutants’ affine angular deviation.(h) Pairwise comparison of MCF10A and PTEN^-/-^ (i) Pairwise comparison of KRas and KRas/PTEN^-/-^. Error bars represent the standard deviation across independent replicates.

We know from analysis of the nuclear tracks that the metastatic MDA-MB-231 cells have a larger average motion than the non-tumorigenic MCF10A cells (**Supplementary Figure S2**). Affine calculations reveal that the MDA-MB-231 cells also display larger deformations and dilations (**Figure 3b**). Thus, the larger average total motion observed in the MDA-MB-231 cells has a collective component. The MCF10A mutants have similar affine dilation and deformation (**Figure 3c**). Combining data from all 10 experiments (**Figure 3d**), the magnitude of affine/collective motion among MCF10A mutants is similar.

Next, we compare the contribution of the two components of motion, affine (collective) and non-affine (individual), to the cell sheet’s direction of motion. Angular distributions of both components (**Figure 3e**) show that the non-affine motion is roughly uniformly distributed in all directions, while the directions of affine motion favor the directional cue provided by empty space in the experimental system (roughly 90 degrees in the figures). This shows that the collective migration of cell sheets is mostly led by affine motion, where individual cells more closely follow their neighbors’ motion. The non-tumorigenic MCF10A and dormant PTEN^-/-^ cell lines show much more directional affine motion than the poorly tumorigenic and metastatic cell lines.

We calculate angular deviation for both affine and non-affine motion among all cell lines to quantify the spread of these distributions. Angular deviation values range from 0 to 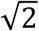, with 0 meaning all cells are moving in one direction and higher values indicating a more dispersed set of migration directions. The non-affine motion of all cell lines has an angular deviation near 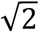, which confirms the observation that non-affine motion has no directional preference (**Figure 3e)**. Meanwhile, the angular deviation differs for the affine (collective) motion across the different cell lines. The affine motion of the MCF10A cells is much more directional than the metastatic MDA-MB-231 cells (**Figure 3f**). Among the MCF10A model system, KRas and KRas/PTEN^-/-^ have significantly higher affine motion angular deviations compared to MCF10A and PTEN^-/-^, indicating the collective motions of the poorly tumorigenic (KRas) and metastatic (KRas/PTEN^-/-^) cell lines are less directional (**Figure 3g**). A pair-wise comparison to MCF10A shows that PTEN^-/-^ cells have the lowest angular deviation (**Figure 3h)**, indicating that PTEN deletion leads to more directional movement consistent with prior PIV analysis [13]. We note that pair-wise comparison of KRas and KRas/PTEN^-/-^ mutants shows that PTEN deletion does not have a similar effect of increasing directionality of collective motion in KRas mutants (**Figure 3i**).

### Quantification of cell rearrangements by D^2^_min_ is robust to tracking errors

To test the robustness of our findings, 20% of the cells were randomly removed in each frame to simulate a cell segmentation error situation, which might occur with non-ideal imaging or segmentation parameters. The U-Net applied here was trained on a different cell line, without any retraining. We manually segmented a set of images and found a segmentation accuracy of at least 80% with respect to detecting all cell nuclei in the sample (**Supplementary Figure S2**). As such, the removal of 20% of the tracked cells is a good simulation to test the measurement robustness of our pipeline to expected segmentation errors. **Figure 4a** shows the original data, with each cell nucleus (segmented by U-Net) labeled in yellow and a frame where 20% of the identified nuclei are labeled for removal (red). Our D^2^_min_ analysis is still able to distinguish between the KRas and KRas/PTEN^-/-^ cell lines, as shown in **Figures 4b, c**. With the removal of 20% of cells, the D^2^_min_ of these four cell lines maintain the same overall trends, including KRas/PTEN^-/-^ consistently exhibiting higher D^2^_min_ than the cell line mutated in KRas alone. The average neighbor distance (**Figure 4d**) slightly increases and is accompanied by a slightly smaller D^2^_min_ for all cell lines with 20% cell removal. So, the larger neighbor distance seen in KRas/PTEN^-/-^ cells cannot explain the higher D^2^_min_ found in these cells. This means that high D^2^_min_ for KRas/PTEN^-/-^ reflects individualistic motion.

**Figure 4:**
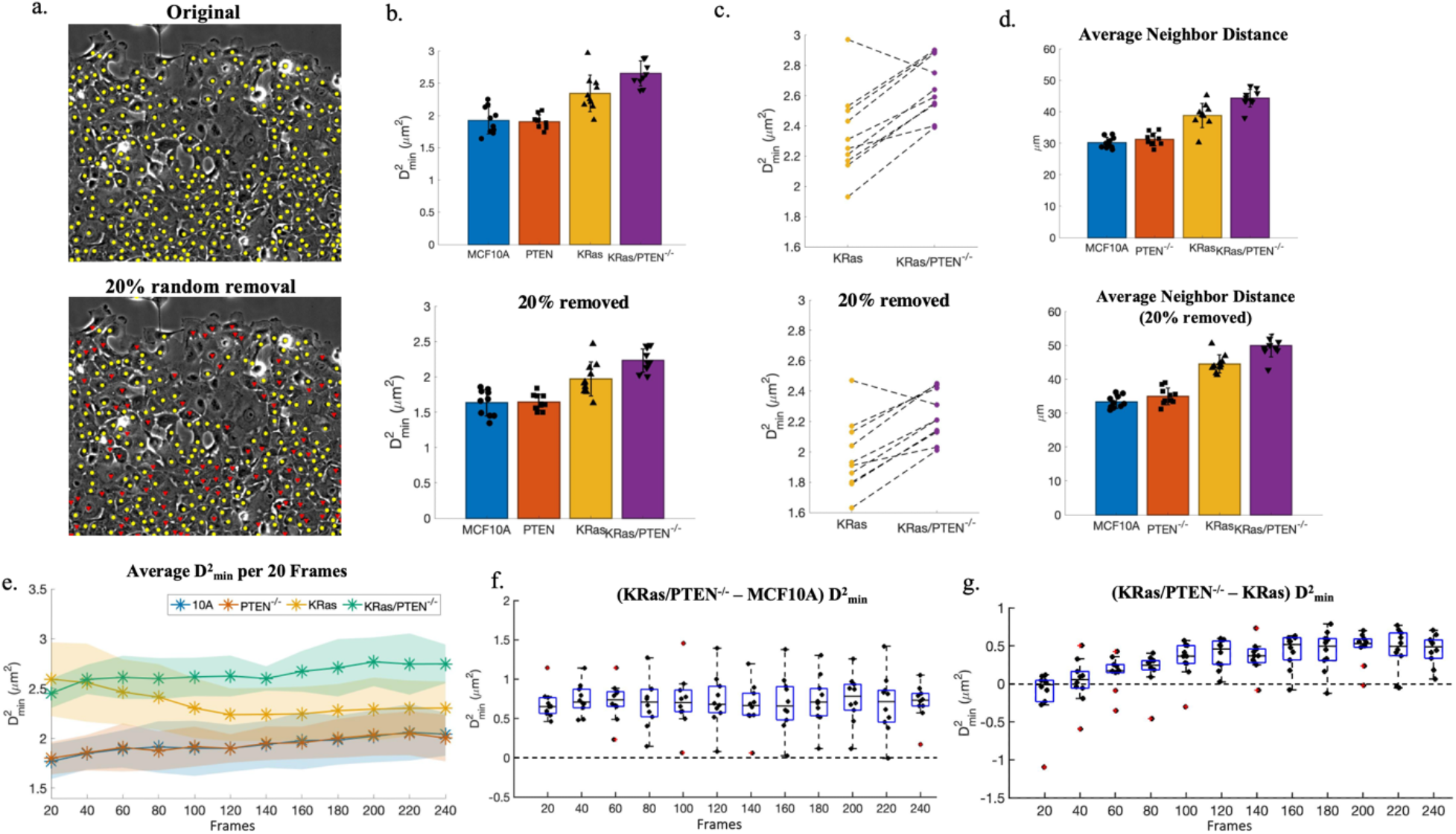
D^2^_min_ is robust to potential tracking errors. (a) Top: a sheet of MCF10A cells, with all U-Net segmented nuclei labeled in yellow; bottom: the same sheet of MCF10A cells, with 20% of nuclei randomly removed from tracking (indicated in red) (b) Top: average D^2^_min_ for 10 independent experiments from U-Net segmented nuclei; bottom: average D^2^_min_ with 20% of nuclei randomly removed in each frame. (c) Pairwise D^2^_min_ comparison of KRas and KRas/PTEN^-/-^ (d) Average neighbor distance. (e) Average D^2^_min_ per 20 frames (60 min) for the MCF10A mutants over time (shaded regions represent the standard deviation across independent replicates). (f) Average D^2^_min_ difference between KRas/PTEN^-/-^ and original MCF10A per 20 frames (60 minutes) over time (g) Average D^2^_min_ difference between KRas/PTEN^-/-^ and KRas per 20 frames (60 minutes) over time. For boxplots, the central line indicates the median value, while the top and bottom edges of the box indicate the 75th and 25th percentiles. Error bars in b, d represent the standard deviation across independent replicates.

Next, we investigate the change in motion with time over the 12 h time scale of our experiments. We segmented the data into 20-frame pieces and found the average D^2^_min_ over each time segment, as shown in **Figure 4e**. The shaded regions in the plot are the standard deviations for D^2^_min_ values over 10 experiments. The poorly tumorigenic KRas and metastatic KRas/PTEN^-/-^ have consistently higher D^2^_min_ compared to the non-tumorigenic MCF10A and dormant PTEN^-/-^ cells at all time points. The KRas cells’ D^2^_min_ value decreases over the first 120 frames, and then stabilizes; the KRas/PTEN^-/-^ mutant D^2^_min_ values slightly increase over time. These two time-related behaviors lead to the KRas/PTEN^-/-^ cells having a higher time-averaged D^2^_min_ value, as shown in **Figures 2e, f**. Over the 12 h of imaging (240 frames), the metastatic KRas/PTEN^-/-^ consistently has higher D^2^_min_ than the non-tumorigenic MCF10A (**Figure 4f**), indicating that D^2^_min_ is a robust comparison between non-tumorigenic and metastatic cells. Comparing the KRas/PTEN^-/-^ and KRas cells (**Figure 4g)**, we find that their motion is initially similar, but differences emerge 120 frames (6 h) after the start of imaging, or 30 h after cell plating. At later times, 20 frames (60 min) of imaging are sufficient for D^2^_min_ to distinguish the KRas/PTEN^-/-^ and KRas cell lines.

## Discussion

We present an AI-enabled label-free nuclear tracking method capable of identifying the motion of cell nuclei and use this information to quantify the collective behavior of panels of cell lines. We study the MCF10A epithelial cell line and its variants with cancer-related mutations, specifically contrasting KRas activation alone and in conjunction with PTEN deletion. These two oncogenic mutations yield different outcomes *in vivo* but their associated motion characteristics have been indistinguishable when analyzed with prior methods [12, 14].

Our nuclear segmentation is done by an AI model previously trained on a human iPSC clonal line [17]. The parameters of the tracking algorithm maintain a temporary memory of cells that disappear in a frame or two due to imaging or segmentation issues and remove tracks that are too short, improving successful cell tracking and reducing noise. With these quality-control features, we can segment and track on average 80% of the nuclei in an image for MCF10A and MDA-MB-231 cell lines without further training on U-Net (**Supplementary Figure S2)**.

The successful application of the AI model on these two cell lines suggests that our AI model and tracking algorithm are versatile and can be applied to other epithelial cell lines without an extra training set. Moreover, random removal of 20% of the nuclei yielded consistent D^2^_min_ results (**Figure 4**), suggesting that we are tracking a large enough fraction of cells for robust calculations not prone to change with tracking errors. Being versatile and robust to errors, we can use our workflow to track and analyze the collective motion for a variety of different types of cells.

Based on measuring the individual motion of cells, D^2^_min_ is capable of clearly distinguishing non-tumorigenic and tumorigenic cells and is also able to distinguish the poorly tumorigenic KRas and the metastatic KRas/PTEN^-/-^ cells (**Figure 2f**). The D^2^_min_ calculation (**Eq 1**), provides multiple metrics for understanding collective migration behavior. In addition to D^2^_min_, which measures the balance of collective vs individual behavior, the calculation quantifies “affine” motion characteristics, namely, the magnitudes and preferred directions of dilations and deformations. Both our previous PIV and current U-Net tracking results show that PTEN deletion makes cells more collective and more directional (**Figure 3)**, while KRas activation makes cells less collective (**Figures 2 and 4)**. However, D^2^_min_ highlights the differences between the poorly tumorigenic KRas and metastatic KRas/PTEN^-/-^ cell lines (**Figure 2**), despite similarities in their collective behavior (**Figure 3**). The metastatic cell line (KRas/PTEN^-/-^) has the most individual motion, which leads to more local rearrangements in the sheet, impacting the overall collective migration of the cell sheet. This individual motion may be the result of a loss in cell adhesion in tumor progression [24], favoring tumor invasion and metastasis. Our analysis of collective and individual motions has the potential to be applied to distinguish non-tumorigenic, tumorigenic, and metastatic cells for easier and faster diagnosis.

In summary, we introduce an AI enabled label-free nucleus tracking workflow for phase contrast imaging that can follow the motion of cells without the need for new AI training data. The comprehensive assessment of cell motion is critical in enabling us to characterize collective motion using D^2^_min_, which provides a local metric of collective motion that shows differences between metastatic and non-metastatic cell lines. Our results indicate that AI enabled collective motion measurements may provide a new perspective on the identification and diagnosis of metastatic risk.

### NIST Disclaimer

Commercial products are identified in this document in order to specify the experimental procedure adequately. Such identification is not intended to imply recommendation or endorsement by the National Institute of Standards and Technology, nor is it intended to imply that the products identified are necessarily the best available for the purpose.

## STAR Methods

### Resource Availability

#### Lead Contact

Further information and requests for resources and reagents should be directed to and will be fulfilled by the Lead Contact, Wolfgang Losert.

#### Materials Availability

This study did not generate new unique reagents.

#### Data and code availability

The code used for analysis in this study has been deposited at GitHub and will be made publicly available upon acceptance for publication. DOIs for the analysis code will be listed in the key resources table. MCF10A and MDA-MB-231 data underlying the figures are available in **Data S1**. The images analyzed in this study are available in the Image Data Resource [25] under DOI [https://doi.org/10.17867/10000163a]. Any additional information required to reanalyze the data reported in this paper is available from the lead contact upon request.

### Experimental Model and Subject Details

#### Cell Culture and Imaging

The images of cell migration used in this work were collected and previously described by Lee, et al. [14, 26]. Briefly, the immortalized human breast epithelial cell lines used were maintained in a humidified atmosphere at 37 °C and 5% CO_2_. Cells were plated in 12 well plates using a complete block design, which allows for pairwise comparisons between cell sheets imaged on the same plate. A 5 μL drop of 1.5 × 10^6^ cells/ml was plated in the center of each well, allowed to adhere for approximately 45 min, and then rinsed to remove non-adherent cells. This created a circular cell sheet, which was then allowed to adhere overnight before imaging. Cells were imaged every 3 min for 12 h using a 10x phase-contrast objective (0.582 μm/pixel) on a microscope with an incubator chamber and a motorized stage for acquiring multiple positions over time. Additional details are available in [14] and the images are available in the Image Data Resource [25] under DOI [https://doi.org/10.17867/10000163a] [26].

### Method Details

#### Image and migration analysis

Cell nuclear segmentation from phase contrast images was performed by a pre-trained U-Net model [17], which generated labeled binary images of inferred connected objects representing nuclei. We used a custom MATLAB code to filter out connected objects that are significantly smaller than the average physiological nucleus size; this step filters out imaging and segmentation noise. The centroid of each nucleus was found using the regionprops function in MATLAB. The Crocker-Grier algorithm [20], as implemented by Blair & Dufresne for MATLAB [27], performs nuclear tracking on the centroids and outputs the tracks of each nucleus for each frame. A custom MATLAB neighbor-finding code [13] was used to find the closest 6 neighbors for each cell nucleus in each frame. This output was then used to calculate D^2^_min_ and the affine components of motion as described in the section below.

Angular deviation, *s*, was calculated as 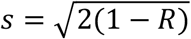, where each tracking vector was transformed into a unit vector in the two-dimensional plane, 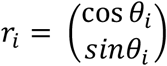, and *R* is the length of the vector average of the direction vectors: 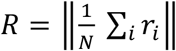

#### D^2^_min_ and affine analysis

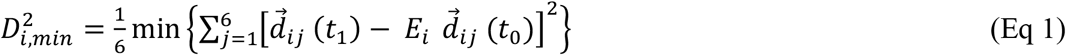

D^2^_min_ is calculated using **Equation 1** by minimizing the difference between the final relative position of a cell and its neighbors,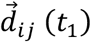, and the affine component, which is a linear transformation of the initial relative position, 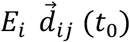.

The 2 × 2 linear transformation matrix *E* that is fit in the D^2^_min_ equation was decomposed to further describe the affine components of motion. We rewrite the matrix as a product of a pure rotation *R*(*θ*) and a symmetric deformation matrix *F* : *E* = *R*(*θ*)*F* which we can expand as shown in **Equation 2**. The angle of rotation, *θ*, can then be found by substituting the elements of *E* into **Equation 3**.

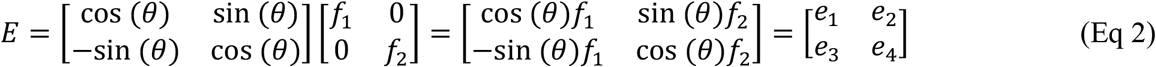

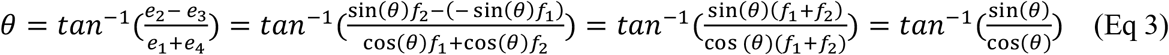

The deformation matrix *F* can then be found using *F* = *R*^−1^(*θ*)*E* − *I*. The subtraction of the identity matrix from *F* ensures that its eigenvalues would be 0 if no deformation, positive if stretching, and negative if compressing. The affine deformation is calculated as the difference between the two eigenvalues of matrix *F*, and affine dilation is the average of the eigenvalues.

The D^2^_min_ calculation is done in the reference frame of an individual cell. Therefore, to extract the affine and non-affine motion components of the lab frame motion, we performed a change of reference frame. For example, we calculated the D^2^_min_ value for a cell with 6 neighbors from a time *t*_*1*_ to *t*_*2*_. Then the expected difference in motion between cell *i* and neighbors *j* at *t*_*2*_ is 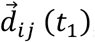, where 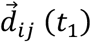 is the difference motion at time 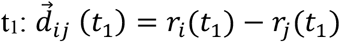. Thus, the expected (affine) neighbor position in the lab frame at time *t*_*2*_ is given by **Equation 4**.

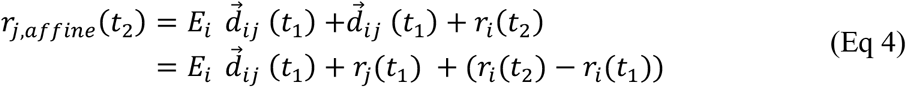

**Equation 4** is the neighbor location in the lab frame plus the cell displacement overtime plus the expected motion difference between cell and neighbors. We then can find the difference between the expected and actual positions for neighbors in the lab frame. Then, taking the average of over 6 neighbors, we get the non-affine motion of the cell compared to its neighbors. To get the affine component of the motion, we simply subtract the calculated non-affine motion from the actual (total) motion.

### Quantification and statistical analysis

For each metric, means shown in the respective figures were calculated from N = 10 independent experiments for the MCF10A genetically defined model system and N = 6 for comparisons to the MDA-MB-231 cells. Error bars represent the standard deviation across independent replicates unless otherwise noted. In boxplot figures, the central line indicates the median value, while the top and bottom edges of the box indicate the 75^th^ and 25^th^ percentiles. Outliers on the boxplots (red plus symbols) were defined as values greater than 1.5 times the interquartile range (IQR indicated by the boxplot whiskers). Underlying data for quantitative figures is presented in tabular format in **Data S1**.

## Funding Acknowledgements

This work was supported by NIH grant R01-CA154624, AFOSR grant FA9550-16-1-0052, American Cancer Society Research Scholar Grant RSG-18-028-01-CSM.

## SUPPLEMENTARY INFORMATION

**Supplementary Video S1**. Tumorigenic mutations change rearrangement in sheets across a genetically defined cancer model system. From left to right: MCF10A, PTEN-/-, KRas and KRas/PTEN-/-cell sheets migrating over 12 hours. Colorful dots represent D^2^_min_ values for individual cells, with their values indicated by the color bar. Scale bars are 100 μm and clocks are shown as HH: MM.

**Supplementary Figure S1:**
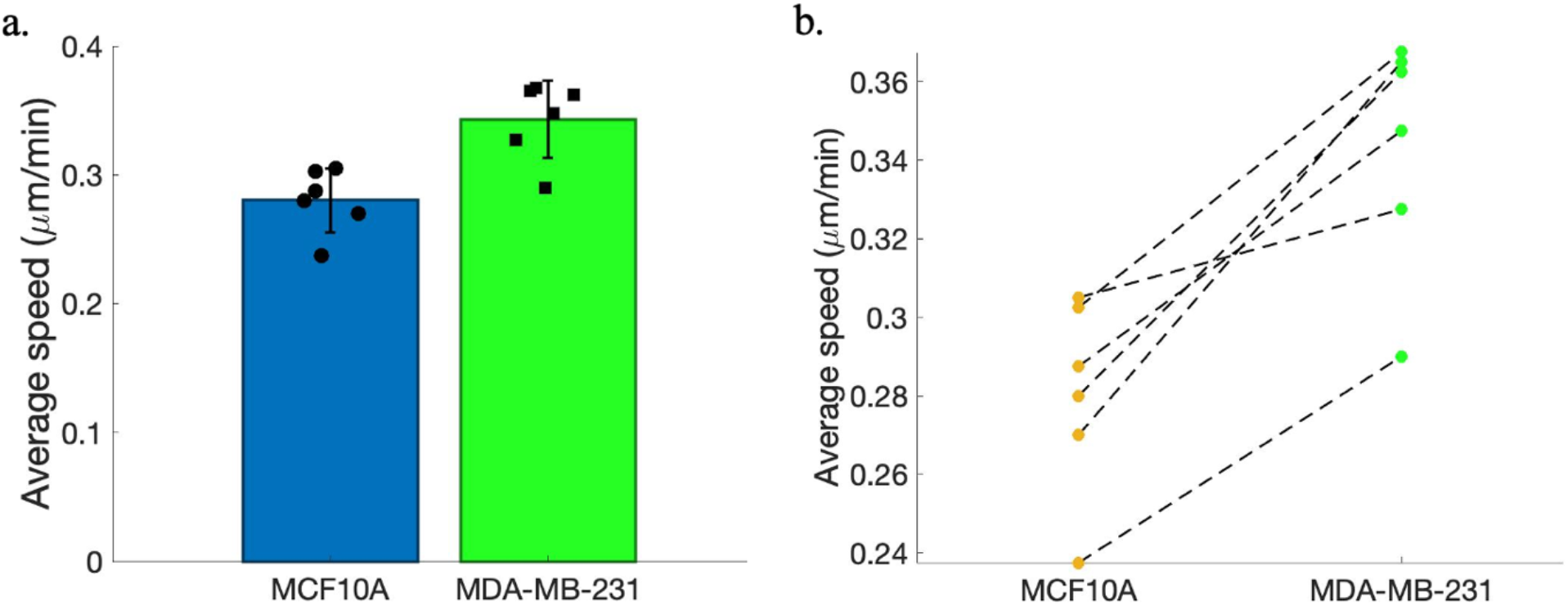
MDA-MB-231 cells exhibit increased overall motion. (a.) The average speed of MCF10A and MDA-MB-231 cells (6 independent experiments). Error bars represent the standard deviation across independent replicates. (b.) Pairwise comparison of MCF10A and MDA-MB-231.

**Supplementary Figure S2:**
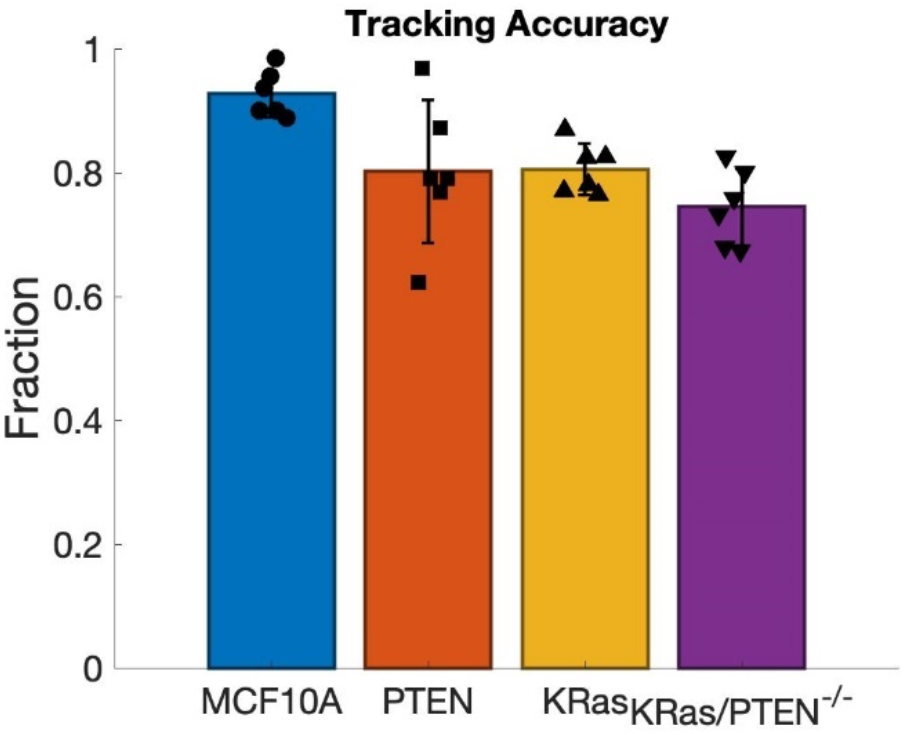
Manual assessment of tracking accuracy. We define accuracy of tracking per frame to be the number of cells found by the tracking minus any double counted cells, divided by the total cell number. The number of cells in each frame were manually counted using the Fiji Cell Counter plugin (https://imagej.nih.gov/ij/plugins/cell-counter.html). We calculated the tracking accuracy on 3 independent experiments for each MCF10A mutants by manually assessing frame 20 (1hour) and frame 220 (11 hour) in the image sequence. Error bars represent the standard deviation.

